# Identification of Candidate Diagnostic gene CDC25C for Trametinib-induced Cardiotoxicity by Integrating Bioinformatics and Machine Learning

**DOI:** 10.1101/2024.05.06.592660

**Authors:** Jixiang Pei, Xiong Yan, Qingming Guo, Zhimei Wang, Yukun Liu

## Abstract

Trametinib is a MEK inhibitor that has been shown to have considerable efficacy in retarding melanoma progression. However, the exact mechanism of cardiotoxicity induced by trametinib remains unclear. Our study was designed to investigate the underlying mechanisms by which trametinib-induced cardiotoxicity (TIC) might exist, offering novel perspectives and guidance for potential prediction, diagnosis, and treatment of TIC. The GSE217421 dataset indicated 574 up-regulated and 705 down-regulated DEGs. According to the KEGG analysis, these genes were implicated in several pathways and functions, including Cell Cycle, Axon Guidance, Cellular Senescence, and Dilated Cardiomyopathy. The GO analysis suggested their association with Mitotic Cell Cycle, Microtubule Cytoskeleton and Adenyl Nucleotide Binding. The hub genes (CDC25C) for TIC were screened through Multiple machine learning algorithms. Next, The expression level of CDC25C was verified using the GSE217423 validation set. Nomogram model based on CDC25C demonstrated excellent diagnostic capability according to three different evaluation measures. To further explore the regulatory mechanism of CDC25C in TIC, we constructed a multi-regulatory network using miRNAs-lncRNAs-TFs-CDC25C and conducted an immunoinfiltration analysis. Our study suggests that CDC25C may be a candidate diagnostic gene and a potential therapeutic target for the early occurrence and development of TIC. This provides new ideas for the prediction, diagnosis and treatment strategies for TIC.

## 1. INTRODUCTION

Melanoma, originating from melanocytes, is a noted type of skin cancer known for its potential virulence.^1^ Although its prevalence only accounts approximately for 1% of skin cancer diagnoses in the United States, melanoma fatalities dominate the mortality statistics for skin cancer. It is estimated that 7,990 deaths (5,420 men and 2,570 women) from melanoma will occur in the United States in 2023.^2^ Positively, from 2011 to 2020, annual melanoma mortality rates exhibited a reduction by roughly 5% in adults below 50, and by about 3% in those aged 50 and above, attributable to advancements in treatment modalities.

Among the promising therapeutic options for melanoma, Trametinib, a MEK inhibitor, has demonstrated considerable efficacy in hindering melanoma progression.^3^ Trametinib’s operational mechanism involves obstructing the proliferation of cancer cells, thereby combating this disease.^4^ However, recent research has shown some deleterious effects associated with Trametinib, notably cardiotoxicity impacting the heart.^5^ Significantly, ∼11% of Trametinib-receiving patients develop a form of cardiomyopathy diagnosed through a decrease in ejection fraction by over 10%, according to echocardiography findings.^7^ The occurrence of cardiotoxicity in patients subjected to Trametinib treatment emerges as a multifactorial process. Studies indicate that signaling via MEK1/ERK1/2 (Extracellular Signal-Regulated Protein Kinase 1/2) is vital for maintaining cardiomyocyte homeostasis and responding to cardiac stress conditions.^6^ However, Trametinib, as a MEK inhibitor, affected normal cardiomyocyte homeostasis through this mechanism, resulting in cardiotoxicity. Trametinib-induced Cardiotoxicity (TIC) results emphasize the necessity of adequately weighing maximum therapeutic advantage against minimum possible harm in clinical practice. Consequently, the identification of risk variables, complemented with the development of superior monitoring and management approaches, becomes indispensable.

At present, with the rise of machine learning, the combination of bioinformatics technology and machine learning algorithms can more effectively solve the problems faced in medicine, as well as better identify the key features required.^8-9^ In our study, the potential candidate diagnostic genes (cell division cycle 25C, CDC25C) for TIC were screened through bioinformatics analysis combined with a variety of machine learning algorithms. After verification by the test set, we established the nomogram diagnosis model for TIC based on CDC25C and tested it by three evaluation models. In order to better understand the role of CDC25C in TIC, we explored it by establishing multi-regulatory networks and immunoinfiltration analysis. Our work provides a viable workflow for exploring the molecular mechanisms of TIC, which can also be used to study cardiotoxicity of other drugs. In conclusion, our study reveals the role of CDC25C in TIC and provides directions and ideas for future research into the potential mechanisms of TIC.

## 2. MATERIALS AND METHODS

### 2.1. Data acquisition

The high throughput sequencing data is downloaded from NCBI GEO databases, including GSE217421 and GSE217423.^10^ GSE217421 serve as test sets, and GSE217423 serves as validation sets. The GSE217421 dataset, the largest for TIC, was derived from the GPL24676 platform and contained trametinib treated and untreated human induced pluripotent stem cells (iPSCs) (19 Tra Group VS 63 Con Group). The GSE217423 dataset was based on the GPL16791 platform and included 12 control samples and 7 trametinib treated samples, also derived from iPSCs.

### 2.2. Identification of different Expression Genes (DEGs)

We used R Bioconductor package limma^11^ for differential expression analysis and identified DEGs. p value was adjusted by Benjamini-Hochberg error discovery rate (FDR), and genes with p value < 0.05 and |Log2 fold-change (log2FC)| > =0.5 were defined as differentially expressed. The volcano map is generated by the R software ggplot2 package.

### 2.3. Enrichment analysis of DEGs

To explore the functions and pathways of common DEGs, we used the R software clusterProfiler package^12^ and the Goplot V1.0.2 package^13^ for functional enrichment analysis (P < 0.05, q < 0.05). For all overlapping DEGs, the term gene ontology (GO) (BP, biological process; CC, cell component; As well as MF(Molecular Function) and Kyoto Encyclopedia of Genes and Genomes (KEGG) pathway enrichment analysis and visualization..

### 2.4. Multiple machine learning algorithms screen for hub genes

Three machine learning algorithms were used to screen for novel and important related genetic biomarkers in TIC: random forest (RF)^14^, Least absolute shrinkage and selection operator (LASSO)^15^, and weighted gene co-expression network (WGCNA)^16^. In this paper, the randomForest R package in R is used to realize the random forest technology. In this study, R software package “glmnet” was used to conduct LASSO logistic regression survey, and the minimum lambda was considered to be optimal. In the next initial phase, the WGCNA R package was used for the construction and modularization of different gene networks at different stages. In our study, genes that share characteristics with more than one of the three classification models discussed previously were selected for further study.

### 2.5. Validation hub gene

In order to eliminate the specificity and clutter of specific data sets, we conducted expression level verification for the validity of hub genes in the GSE217423 dataset, and directly displayed the expression levels of pivot genes between trametinib treated group and control group in the form of violin and box graphs. In addition, we also detected the expression levels of hub genes in GSE217421 test sets. In short, the hub genes that were co-expressed in the three datasets were involved in the subsequent analysis.

### 2.6. The diagnostic and prognostic model was established and evaluated based on candidate diagnostic genes

To evaluate the diagnostic value of hub genes for TIC, we evaluated the diagnostic performance of key genes and nomograms by calculating the area under the ROC curve.^17^ We then used ROC curves to evaluate the effectiveness of nomogram based decision making for the diagnosis of TIC. To fully evaluate the predictive accuracy and clinical application of nomogram in TIC, we performed calibration curve and decision curve analysis (DCA).

### 2.7. Multi-factor regulatory network of key diagnostic markers

To understand the factors influencing the regulation of key biomarkers, we used RNAInter,^18^ hTFtarget,^19^ and mirDIP^20^ to predict the involvement of long non-coding Rnas (lncRNAs), microRNAs (miRNAs), and transcription factors (TF) with biomarkers. The multifactor regulatory network was then visualized using Cytoscape software.

### 2.8. Immunoinfiltration analysis

We used the CIBERSORT algorithm^21^ to assess immune cell infiltration in trametinib induced cardiotoxicity and generate a correlated heat map. In addition, we explored differences in immune cell profiles between different groups and analyzed correlations between key biomarkers and immune cells.

## 3. RESULTS

### 3.1. Identification of different expression genes

The high-throughput sequencing expression profile GSE217421 was downloaded from GEO database and analyzed by the limma package of R language. Different expression genes (DEGs) of GSE217421 were screened with false discovery rates (FDR) < 0.05 and |Log2 fold-change (log2FC)| > =0.5 as the screening conditions (FIG. 2A). The heat map shows the top 100 DEGs in each sample and shows their expression in different sample sets (FIG. 2B). In the GSE217421 dataset, the number of DEGs up-regulated and down-regulated were 574 and 705, respectively (FIG. 2C).

**FIG. 2:**
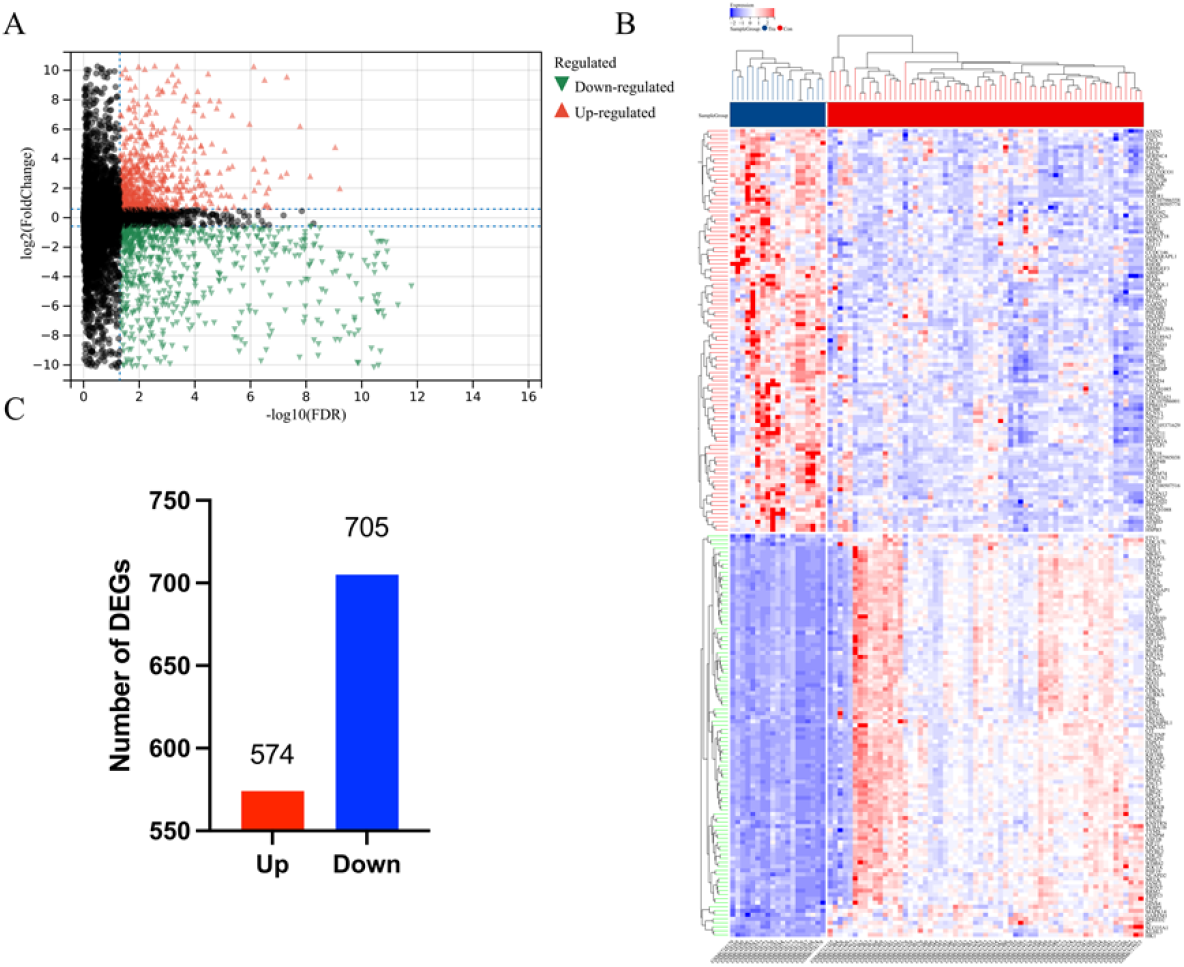
Identification of DEGs. (A) volcano plot; (B) heat map; (C) The number of DEGs.

### 3.2. Functional enrichment analysis

In order to further understand the functions and pathways involved in DEGs, we further enriched them from Kyoto Encyclopedia of Genes and Genomes (KEGG) and Gene ontology (GO). KEGG results showed (FIG. 3A) that DEGs was mainly enriched in the Cell cycle [-log10(P.adjust) =13.04], Axon guidance [-log10(P.adjust) =1.32], Cellular senescence [-log10(P.adjust) =3.47] and dilated cardiomyopathy [-log10(P.adjust) =2.77]. GO analysis results showed (FIG. 3B-D) that DEGs was mainly involved in Cell cycle [-log10(P.adjust) =38.76], Cell cycle process [-log10(P.adjust) =38.77], Mitotic cell cycle [-log10(P.adjust) =39.17] and Protein containing complex subunit organization [-log10(P.adjust) =6.68] in BP; Chromosome [-log10(P.adjust) =5.90], Microtubule cytoskeleton [-log10(P.adjust) =10.11], Supramolecular complex [-log10(P.adjust) =8.37] and Catalytic complex [-log10(P.adjust) =1.51] in CC; Enzyme binding [-log10(P.adjust) =9.59], Ribonucleotide binding [-log10(P.adjust) =10.33], Adenyl nucleotide binding [-log10(P.adjust) =11.42] and Identical protein binding [-log10(P.adjust) =4.28] in MF.

**FIG. 3:**
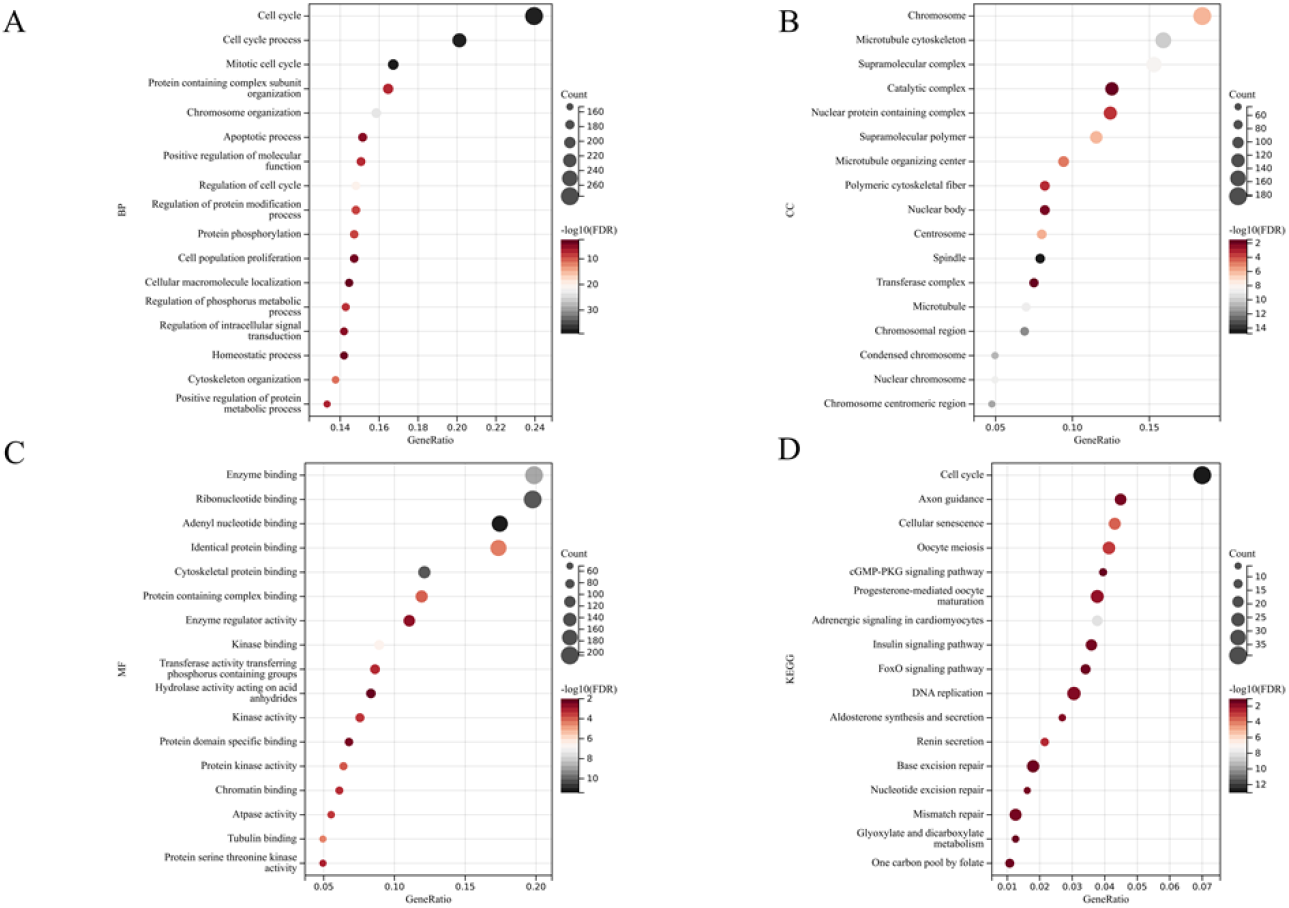
Functional enrichment analysis (A) KEGG pathway; (B) Biological processes (BP); (C) Cell component (CC); (D) Molecular function (MF).

### 3.3. Multiple machine learning algorithms screen for hub genes

To screen for the best hub genes, we integrated several algorithms, including WGCNA analysis, LASSO logistic regression and RF algorithm. WGCNA analysis (FIG. 4A-B) revealed the presence of 27 significant co-expression modules (FIG. 4C-D). The results of module analysis showed that there was an association between multiple modules and the trametinib treatment group (FIG. 4G). Among them, florawhite module was the most significant, resulting in the screening of 143 genes. In RF analysis (FIG. 4G-H), the score rankings of top 10 genes were obtained according to the MeanDecreaseGini score (FIG. 4I). In addition, LASSO multifactor survival analyses were performed on DEGs to determine the optimal number of screening traits (FIG. 4J), and 20 genes were identified (FIG. 4K). Finally, through Venn diagram analysis (FIG. 4O), the intersection of the 3 algorithms is CDC25C.

**FIG. 4:**
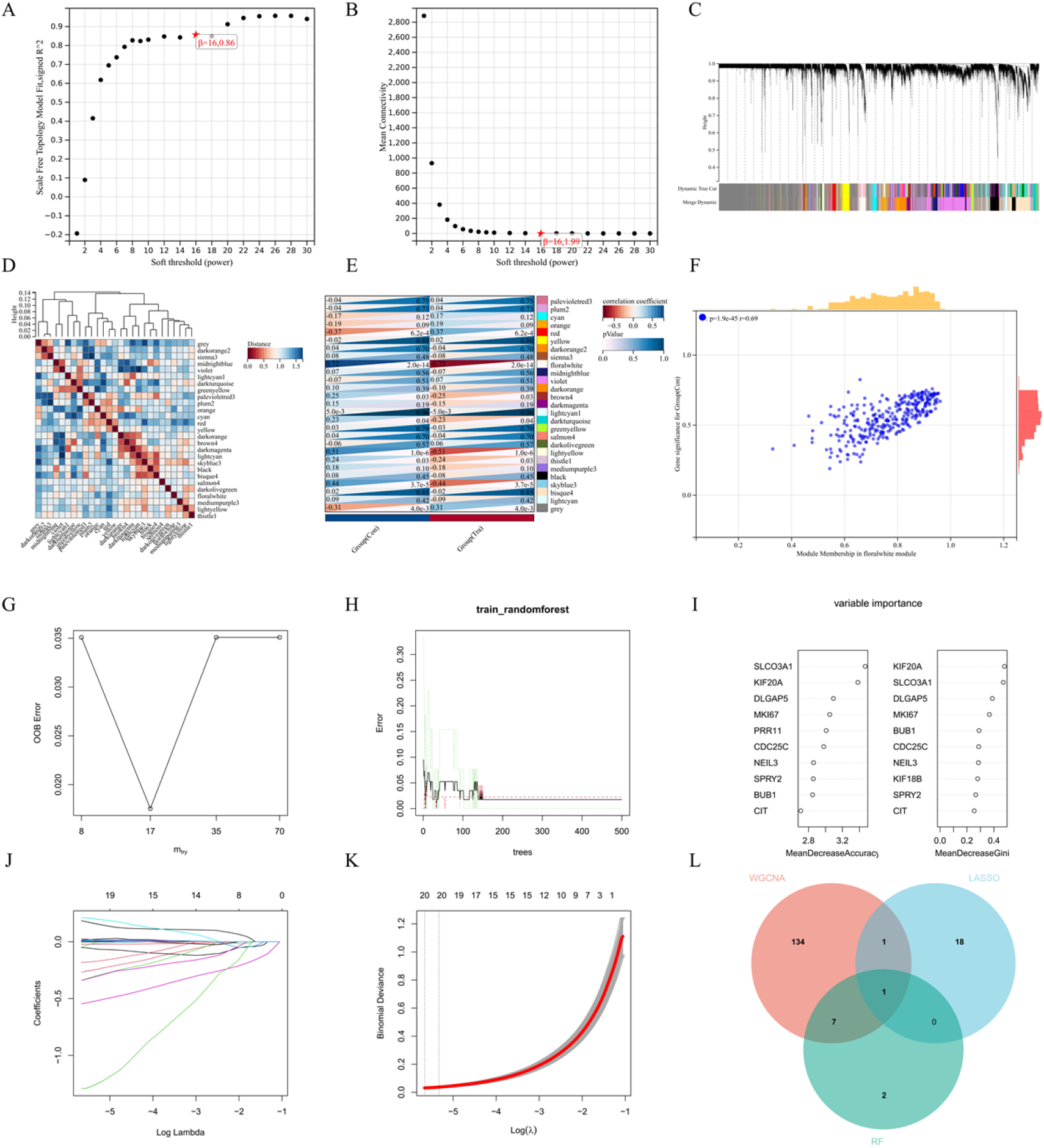
Multiple machine learning algorithms screen for hub genes (A) Average connectivity analyzed by WGCNA; (B) WGCNA analytical scale independence; (C) WGCNA gene clustering; (D) WGCNA module feature vector clustering; (E) Heat map of correlation between WGCNA module and phenotype; (F) GS-MM correlation scatter plot of florawhite module; (G-H) Random forests; (I) MeanDecreease Accuracy and MeanDecreeaseGini score; (J) Multifactor survival analysis; (K) K-M curve; (L) Three algorithmic Venn diagrams.

### 3.4. Validation of CDC25C and Construct diagnostic models

To evaluate the diagnostic value of CDC25C in TIC, ROC curves were established based on two test sets, GSE217421 and GSE217423, and the expression levels of CDC25C were evaluated. Analysis of the GSE217421 dataset showed that the expression levels of CDC25C in the trametinib group were significantly lower than those in the control group (FIG. 5A). Moreover, CDC25C showed good diagnostic ability, with ROC curve values of 0.9833, respectively (FIG. 5B). In order to further verify the effectiveness of screening genes and eliminate the differences and miscellaneous among different data sets, we used the GSE217423 dataset as an external validation set. As shown in FIG. 5C, the expression curves of CDC25C in GSE217423 were consistent with the GSE217421 dataset, and the ROC curves of CDC25C was 0.9762 (FIG. 5D). According to the above analysis, CDC25C may play a key role in the TIC. Therefore, we constructed a diagnostic and prognostic model based on CDC25C and evaluated it. By using logistic regression analysis, we constructed a Nomogram model based on CDC25C. FIG. 7D shows that as CDC25C expression levels decreased, the total score increased, and the risk of cardiotoxicity increased significantly. To test the validity of the Nomogram model, we constructed calibration curves, DCA decision curves and ROC curves to evaluate the model. The ROC curve also proved the diagnostic capability of the Nomogram model, with a value of 0.933 (FIG. 6C). Figure 7G-H shows that the constructed Nomogram model has good diagnostic capability.

**FIG. 5:**
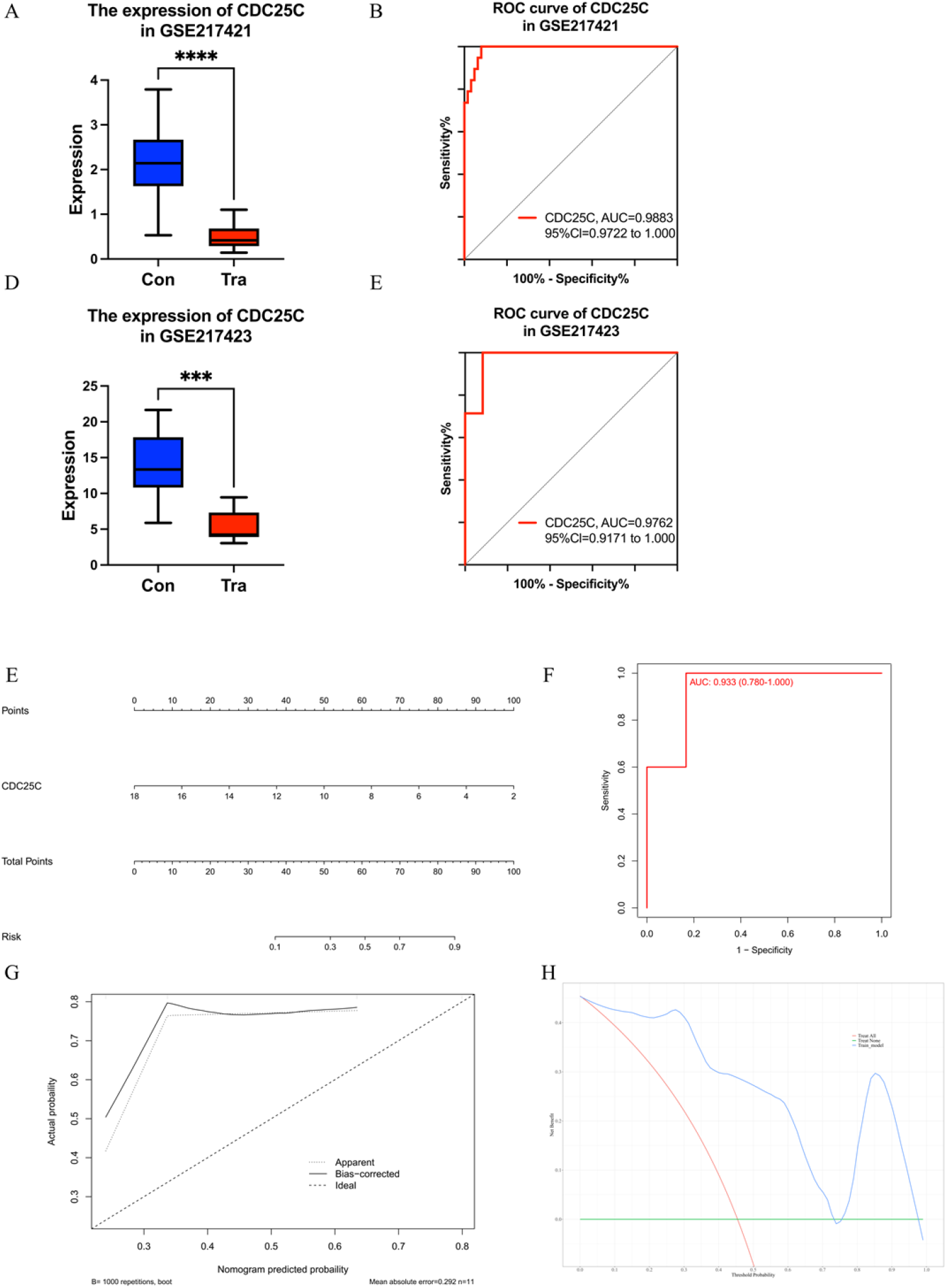
Validation of CDC25C and Construct diagnostic models (A) The expression levels of CDC25C in GSE217421 test set; (B) ROC curves of CDC25C in the GSE217421 test set; (C) The expression levels of CDC25C in the GSE217423 validation set; (D) ROC curves of CDC25C in the GSE217423 validation set; (E) Nomogram model; (F) ROC curve of Nomogram model; (G) Calibration curves; (H) DCA decision curve. (*** means p < 0.001; **** means p < 0.0001)

**FIG. 6:**
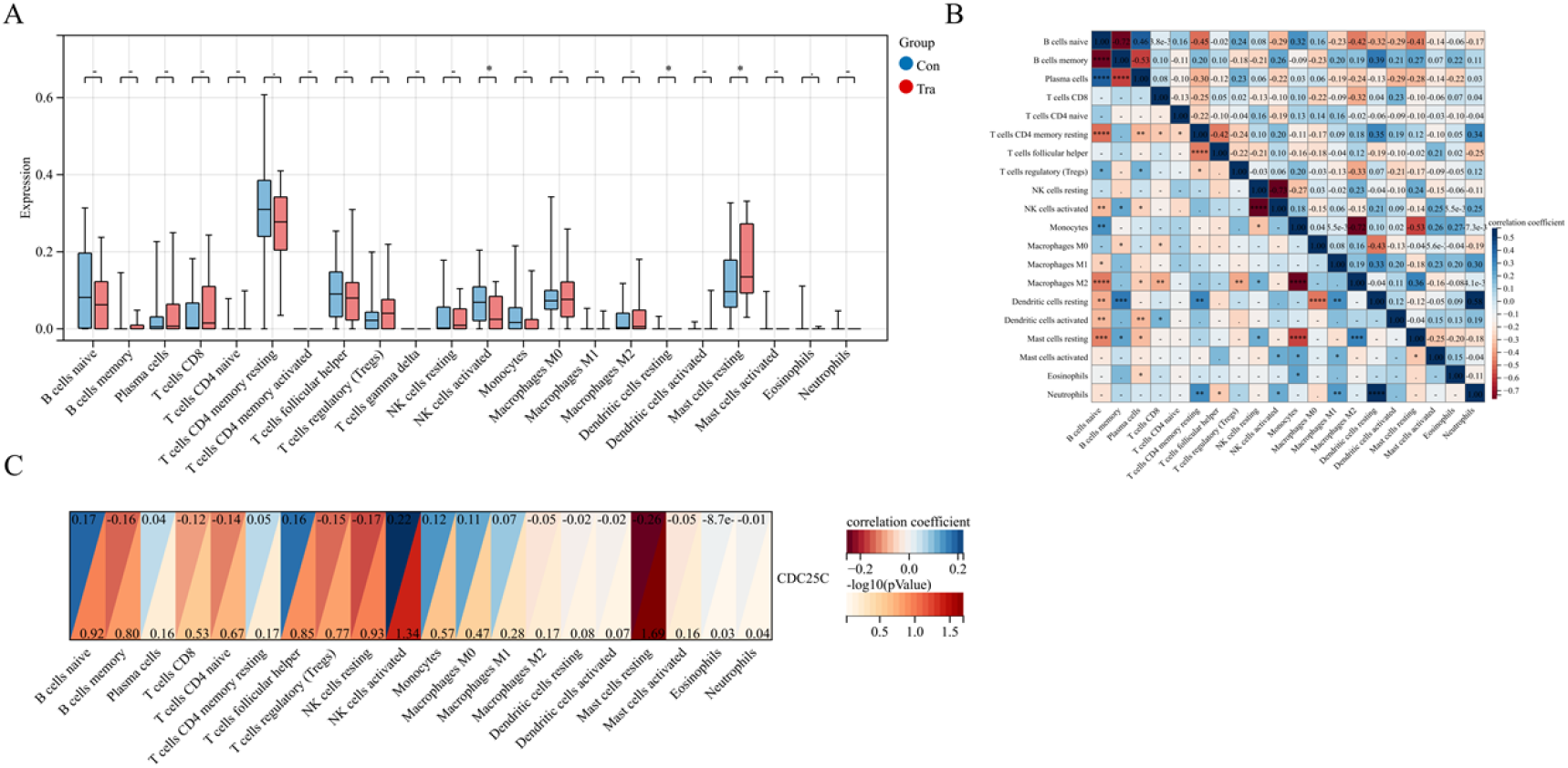
Immunoinfiltration analysis (A) The expression levels of 22 kinds of immune cells; (B) The correlation of 22 immune cellss; (C) Correlation between CDC25C and immune cells. (-means p < 0.05; * means p < 0.05)

**FIG. 7:**
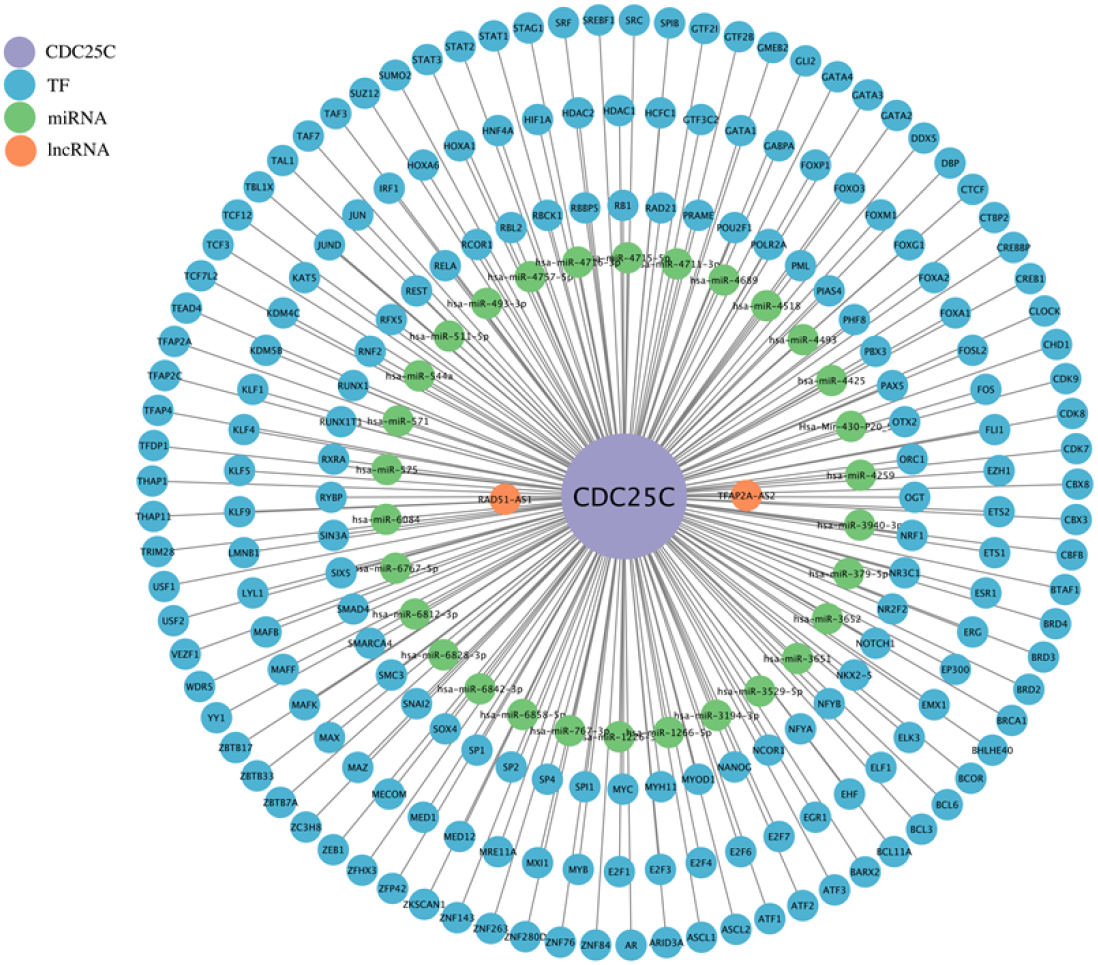
Multi-regulatory network network. The colors of the different circles represent different factors: purple is lncRNA, green is micRNA, blue is TF, and the central circle is CDC25C.

### 3.5. Immunoinfiltration analysis

Previous studies have shown that immune cells play a crucial role in cardiotoxicity. To explore the role of immune cells in TIC, we evaluated the GSE21742 dataset using the CIBERSORT algorithm. FIG. 6A shows that of the 22 immune cells identified, only NK cells activated, dendritic cells resting and mast cells resting show differences between two groups. Moreover, NK cells activated and Dendritic cells resting were significantly reduced in the trametinib treatment group. Compared to the control group, the trametinib treatment group showed higher expression of Mast cells resting. In the correlation heat map (FIG. 6B), NK cells activated showed the most negative correlation with NK cells resting, while NK cells activated showed the most positive correlation with B cells memory. Similarly, Dendritic cells resting were most negatively correlated with Macrophages M0, and Dendritic cells resting were most positively correlated with B cells memory. Then, Mast cells resting showed the most negative correlation with Monocytes, while Mast cells resting showed the most positive correlation with Macrophages M2. CDC25C is a key marker of TIC, and the relationship between CDC25C and immune cells was studied. FIG. 6C showed that CDC25C was most positively correlated with NK cells activated [-log (pvalue) = 1.34], and SLC6A6 was most negatively correlated with Mast cells restings [-log (pvalue) = 169].

### 3.6. Construction of multi-regulatory network

In order to further explore the potential mechanism of CDC25C, we established the regulatory network of lncRNA/micRNA/ TF-CDC25C. The results showed that CDC25C had regulatory relationships with 31miRNAs, 2 lncRNAs and 63 TFs. The colors of the different circles represent different factors: orange is lncRNA, green is miRNA, blue is TF, and the central circle is CDC25C. Our future studies will focus on these regulatory factors and explore the important role of the regulatory relationship between these factors and CDC25C in TIC.

## 4. DISCUSSION

With more than 1 million Americans currently living with melanoma, MEK1 inhibitors have significantly improved patient survival and prognosis.^2-3^ Despite the significant efficacy, the long-term use of MEK1 inhibitors is limited by cardiovascular toxicity. The exact mechanism of TIC remains elusive, posing challenges in predicting or preventing adverse events in patients. It is very necessary to explore effective biomarkers of TIC, which can provide new ideas and insights for the diagnosis, treatment and prediction of chemotherapy-induced cardiotoxicity in the future.

In this study, we identified DEGs in GSE217421, in order to explore important genes involved in TIC. Then, We conducted annotations of these DEGs using KEGG and GO respectively, to gain a deeper understanding of the potential pathways and functions these genes might be implicated in. The findings suggest that the DEGs could be involved in several pathways and functions: Cell cycle, Axon guidance, Cellular senescence, and dilated cardiomyopathy according to KEGG analysis, while GO analysis points towards Mitotic cell cycle in BP, Microtubule cytoskeleton in CC, Adenyl nucleotide binding in MF. Next, in order to select the suitable genes, we utilized a combination of different algorithms, including WGCNA analysis, LASSO logistic regression, and RF algorithm. Initially, the WGCNA results pointed to ‘florawhite’ as the module having the strongest correlation with the phenotype and included 143 genes. Concurrently, the LASSO multifactor survival penalty analysis provided a shortlist of 20 genes from the DEGs. The RF analysis presented us with the top 10 genes according to the MeanDecreaseGini score. Ultimately, the common intersection of these three methodologies was CDC25C, as visualized through a Venn diagram. To mitigate the risk of false positives within our analysis, we performed an expression validation on GSE217423 validation sets. Primarily, we observed that the expression levels of the hub gene CDC25C remained consistent in the two samples, both demonstrating good diagnostic proficiency. Subsequently, we constructed a Nomogram diagnostic model predicated on the expression level of CDC25C to predict TIC. The performance of this model was appraised utilizing the ROC curve, calibration curve, and DCA curve. The results of these evaluations confirm the diagnostic capabilities of our nomogram model.

The Cell Division Cycle-25C dual-specificity phosphatase (CDC25C), which plays a pivotal part in managing serine/threonine kinase function within the cell cycle, can stimulate the G2/M switch in mitotic cells via cyclin-dependent kinase-1 (CDK1) dephosphorylation, thereby triggering the cyclin B1/CDK1 complex.^22^ A gamut of studies indicates that CDC25C manifests itself in various organs such as the prostate,^23^ lung,^24^ bladder,^25^ stomach,^26^ in addition to specific cancers like esophageal colon cancer,^27^ acute myeloid leukemia,^28^ and squamous cell carcinoma.^29^ As in-depth research on CDC25C continues, scientists have come to appreciate its significant potential as a therapeutic target for next-generation cancer treatments, given that some patients experiencing tumors exhibit CDC25C’s immunogenicity, earning it the status of a new tumor marker.^30^ Regrettably, literature pertaining to CDC25C’s specific role in cardiotoxicity is scarce, creating the need for studies like ours, which delves into the critical part CDC25C plays in TIC through the lens of bioinformatics and machine learning, scaffolding multi-regulatory networks and immune infiltration relationships to decipher the potential machinations of CDC25C.

At present, a number of studies have revealed the important role of miRNAs, lncRNAs and TFs in cardiovascular diseases. It is reported that microRNAs can regulate the expression of cyclins CDC25C.^31-32^ miR-125b has been proved to play an anti-apoptotic role by inhibiting apoptosis regulators such as TP53 and BAK1. Overexpression of miR-125b repressed the endogenous level of P53 protein and suppressed apoptosis by regulating the expression of cyclin C and CDC25C.^33-34^ In our research, to further explore the regulatory mechanism of CDC25C in TIC, we established a multi-regulatory network of miRNAs-lncRNAs-TFs-CDC25C. We found that CDC25C had regulatory relationships with 31 miRNAs, 2 lncRNAs and 63 TFs. In the future, we will further analyze these regulatory relationships and explore the possible regulatory mechanisms. Our study also explored the role of immune cells in TIC and the association of immune cells with CDC25C. Immunoinfiltration analysis showed NK cells activated and Dendritic cells resting were significantly reduced in the trametinib treatment group. Compared to the control group, the trametinib treatment group showed higher expression of Mast cells resting. Further analysis found that CDC25C was most positively correlated with NK cells activated [-log (pvalue) = 1.34], and SLC6A6 was most negatively correlated with Mast cells restings [-log (pvalue) = 169].

## 5. CONCLUSIONS

Our study revealed that the expression difference of CDC25C is related to TIC, and its potential mechanism may be related to immune cell infiltration and lncRNA/miRNA/TF regulation mechanism. CDC25C may be a candidate diagnostic gene and a potential therapeutic target for the early occurrence and development of TIC, providing new ideas and directions for the prediction, diagnosis and treatment of TIC in the future, but its diagnostic ability still needs to be further verified.

## ACKNOWLEDGMENTS

The present study was supported by Chen Xiao-ping Foundation for the Development of Science and Techonology of Hubei Province (No.CXPJJH122011-011), Wu Jieping medical foundation (No.320.6750.2023-18-21), Qingdao Medical and Health Research Project (No. 2021-WJZD090)..

## ABBREVIATION

TIC: Trametinib-induced cardiotoxicity
NCBI: National Center for Biotechnology Information
GEO: Gene Expression Omnibus
DEGs: Differentially expressed genes
GO: Gene ontology
KEGG: Kyoto Encyclopedia of Genes and Genomes
WGCNA: Weighted gene co-expression network
RF: Random forests
LASSO: Least absolute shrinkage and selection operator
ROC: Receiver operating characteristic
AUC: Area under the curve
DCA: Decision curve analysis
PPI: Protein-protein interaction
CIBERSORT: Cell-type Identification By Estimating Relative Subsets Of RNA Transcripts
lncRNAs: Long non-coding RNAs
miRNAs: MicroRNAs
TFs: Transcription factors
CDC25C: Cell division cycle 25C

## Ethics approval and consent to participate

Not applicable.

## Consent for publication

All authors gave their consent for publication.

## Availability of data and materials

The datasets used and/or analyzed during the current study are available from the corresponding author on reasonable request. The data that support the results of current study is available on Gene Expression Omnibus (GEO).

